# Palmitoylation of PSD95 flips the bilayer juxtaposed domain and drives cluster formation on membrane

**DOI:** 10.1101/2025.10.09.679381

**Authors:** Pritakshi Das, Deepak Nair, Anand Srivastava

## Abstract

Postsynaptic density protein 95 (PSD95) is a highly abundant scaffolding protein at excitatory synapses, where it organizes the molecular architecture of the postsynaptic density (PSD) by anchoring neurotransmitter receptors, ion channels, adhesion molecules, and signaling enzymes. Emerging evidence indicates that PSD95 forms transient “nanodomains” essential for the fidelity of synaptic transmission and plasticity, and disruptions in its clustering are linked to neuropsychiatric disorders. Palmitoylation, a reversible post-translational modification involving the covalent attachment of palmitate to Cysteine residues, has been implicated in the localization and clustering of the PSD95 at the plasma membrane, but its structural and mechanistic role remains unclear. Here, we use a multiscale molecular dynamics approach that combines all-atom and coarse-grained simulations to investigate how palmitoylation regulates PSD95 conformational dynamics, membrane association, and self-assembly. Our results show that palmitoylation promotes an open conformation in solution and stabilizes an extended assembly-primed state at the membrane. In the unmodified state, the PDZ1 domain remains distal while PDZ2 preferentially engages with the membrane; palmitoylation reverses this arrangement, anchoring PDZ1 to the membrane and enhancing clustering propensity. Membrane lipid composition further modulates this behavior, with negatively charged lipids influencing favorable domains orientation through electrostatic interactions. Finally, coarse-grained simulations reveal that palmitoylated PSD95 forms stable dimers on the membrane surface, whereas unmodified molecules fail to associate persistently. Together, these findings provide a mechanistic model for how palmitoylation facilitates the early stages of PSD95 clustering, offering nanoscale insights into synaptic organization that complement experimental observations.

**Significance Statement:** PSD95 is a key scaffolding protein that organizes synaptic components essential for neuronal communication. Its ability to cluster at the membrane is critical for postsynaptic architecture, yet the molecular determinants of this process remain unclear. Using multiscale molecular dynamics simulations we reveal how palmitoylation, a reversible lipid modification, acts as a molecular switch that anchors PSD95 to the membrane and reorients its domains to favor self-association. We further demonstrate that membrane lipid composition modulates these effects through electrostatic interactions. Our work offers a mechanistic model for the initial steps of PSD95 clustering, highlighting how post-translational modifications and membrane environment together regulate synaptic protein organization. These insights bridge molecular structure with synaptic function and are not easily accessible through current experimental methods.

**Graphical Abstract Image:** 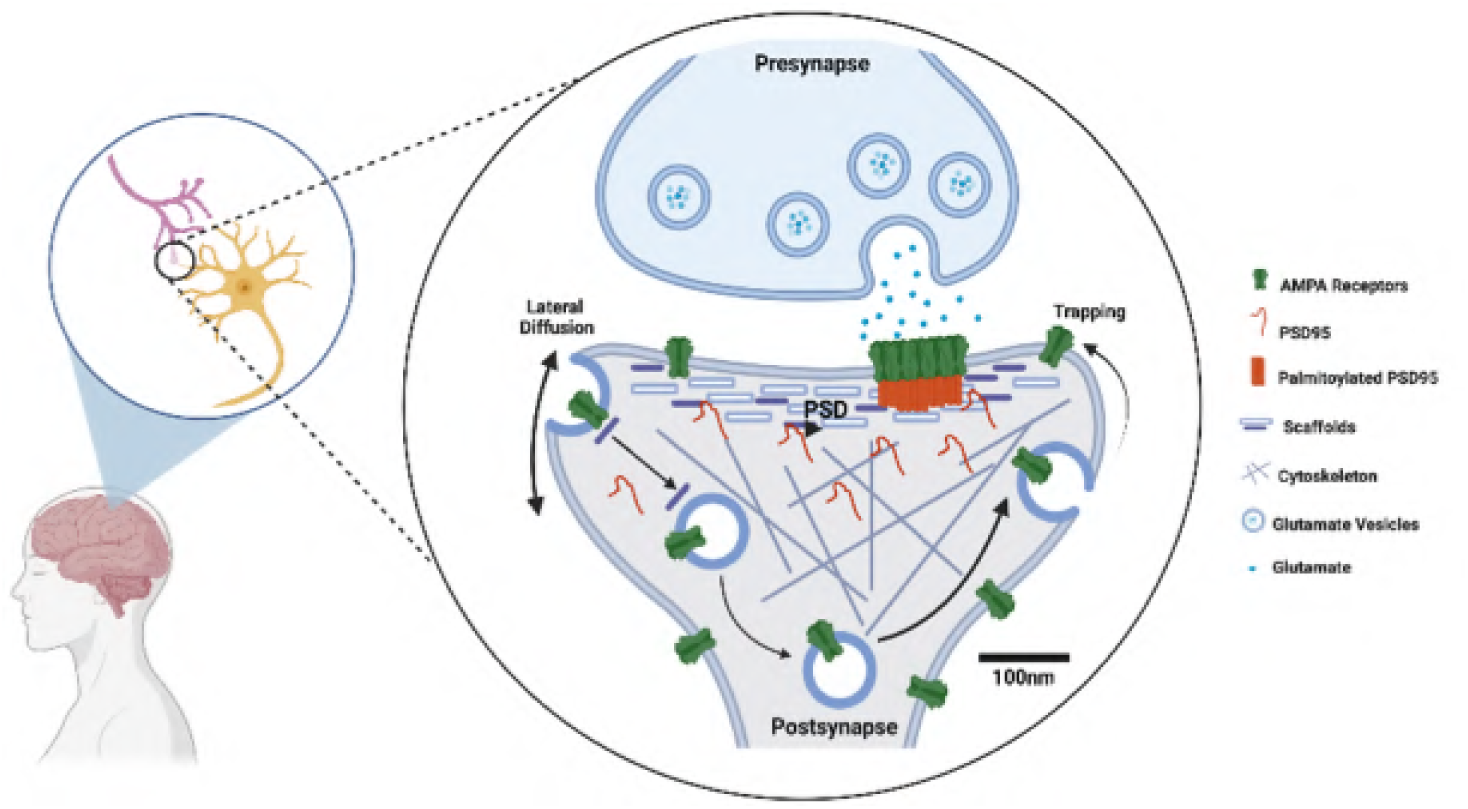

## 1 Introduction

Membrane-associated guanylate kinases (MAGUKs) are multidomain scaffold proteins that play a pivotal role in organizing signaling complexes at the membrane, particularly at the postsynaptic density of neurochemical synapses and other cell junctions.^1–3^ Among MAGUKs, the Discs large (DLG) subfamily is crucial for the molecular organization of cell adhesion complexes in neurons and epithelial cells. This subfamily includes SAP97/hDLG, SAP90/PSD95, SAP102/NE-DLG, and PSD93/Chapsyn 110, all of which share a unique modular structure composed of several protein–protein interaction domains. ^4–6^ To date, all known members of the DLG family contain three PDZ (PSD95/DLG/ZO1) domains, an Src homology 3 (SH3) domain, and a guanylate kinase (GUK) domain. Among these, the PDZ domains are the best characterized domains, exhibiting high-affinity binding to carboxyl-terminal peptide motifs of various proteins, such as GluN2 subunits of NMDA receptors, GluA1 subunits of AMPA receptors, and voltage-gated inwardly rectifying Potassium (K+) channels.^7^ PDZ domains also bind to many cell ashesion molecules and also to TARPS-like stargazin proteins.

Interestingly, the GUK domain of MAGUKs lacks the essential amino acids required for ATP or GMP binding, indicating it no longer serves an enzymatic function. Instead, it mediates protein–protein interactions, such as those with multidomain proteins like GKAP and SPAR.^8,9^ The SH3 domain also exhibits an unusual binding specificity, capable of both intra- and intermolecular interactions with the GUK domain. Intramolecular binding is thought to be more prevalent and is stabilized by additional tertiary interactions when the SH3 and GUK domains are in close proximity on the same polypeptide chain.^8,10^

Among DLG family members, postsynaptic density protein 95 (PSD95) is a key player in the organization and function of excitatory synapses in the brain. As a multi-domain scaffolding protein, PSD95 acts as an anchor, tethering a variety of neurotransmitter receptors and signaling molecules to the postsynaptic membrane, thereby facilitating synaptic maturation and plasticity.^11^ Understanding the mechanisms that govern PSD95 clustering is essential for elucidating synaptic function. One critical regulatory process is palmitoylation, a post-translational modification involving the attachment of a fatty acid group, palmitate, to Cysteine residues 3 and 5 of PSD95.^12–15^ Palmotylation motifs are nor present in every PSD95 however large fraction of PSD95 exists in the plamotylated isoform or alpha isoform. Palmitoylation enhances PSD95’s membrane association and promotes the formation of stable nanodomains at the postsynaptic density, which serve as hubs for organizing signaling complexes and ensuring effective synaptic transmission.^16–21^

A key interaction in this context involves stargazin, a member of the transmembrane AMPA receptor regulatory proteins (TARPs), which plays a crucial role at the synapse. Stargazin enhances the synaptic localization of AMPA receptors and facilitates their clustering by directly binding to the PDZ domains of PSD95.^22–25^ This interaction is mediated by the carboxy-terminal PDZ-binding motif of stargazin, which docks specifically into the PDZ1 and PDZ2 domains of PSD95 with high specificity.^23^ The structural complementarity between stargazin’s terminal sequence and the PDZ domains of PSD95 ensures the formation of a stable, high-affinity complex, anchoring AMPA receptors to the postsynaptic density and stabilizing their membrane localization for efficient synaptic transmission.^23,26^ Stargazin’s interaction with PSD95 is dynamically regulated by phosphorylation and the lipid environment of the membrane. Phosphorylation of stargazin’s intracellular C-terminal tail by kinases such as calcium/calmodulin-dependent protein kinase II (CaMKII) modulates its affinity for PSD95, thereby tuning the recruitment and retention of AMPA receptors at synapses. This activity-dependent regulatory mechanism underlies the remodeling of AMPA receptor localization during synaptic plasticity, including long-term potentiation (LTP) and long-term depression (LTD).^27–29^

The formation of PSD95 nanodomains facilitates cooperative interactions between stargazin and other PSD95 binding partners, such as additional TARPs or GluA subunits of AMPA receptors. This multivalent interaction network, stabilized by PSD95’s palmitoylation and modular domain structure, is essential for the precise spatial arrangement of postsynaptic signaling complexes. The interplay between PSD95 and stargazin exemplifies a highly coordinated mechanism of synaptic organization, where post-translational modifications, domain-specific interactions, and membrane localization collectively regulate receptor clustering and synaptic efficacy. By promoting the formation of stable PSD95 nanodomains, palmitoylation not only strengthens stargazin-PSD95 binding but also ensures the dynamic adaptability of synapses to changing neuronal activity.

However, the precise molecular mechanisms by which palmitoylation regulates PSD95 clustering remain to be fully elucidated. To address this, our study employs multiscale molecular dynamics (MD) simulations to investigate how palmitoylation influences PSD95’s conformation and interactions with the membrane. Fig. 1 shows the domain organization and structural modeling approach we used. The five distinct domains of PSD95 are connected by flexible linker regions, allowing the protein to adopt a wide range of conformations. Notably, palmitoylation occurs at Cysteine residues 3 and 5, which plays a key role in membrane association and protein clustering (Fig. 1a). We reconstructed the full-length PSD95 by integrating structural predictions from I-TASSER with a comprehensive analysis of the model (Fig. 1b), this initial model was used for all molecular dynamics simulations performed in this study, providing insights into PSD95’s conformational dynamics and interactions (Fig. 1c). These simulations reveal that palmitoylation induces specific conformational changes in PSD95, facilitating its membrane association and enhancing interactions between individual PSD95 molecules. This ultimately promotes the formation of robust PSD95 clusters. Investigating these processes provides new insights into synaptic regulation and potential implications for neurodevelopmental disorders linked to PSD95 dysfunction.^30^

**Figure 1:**
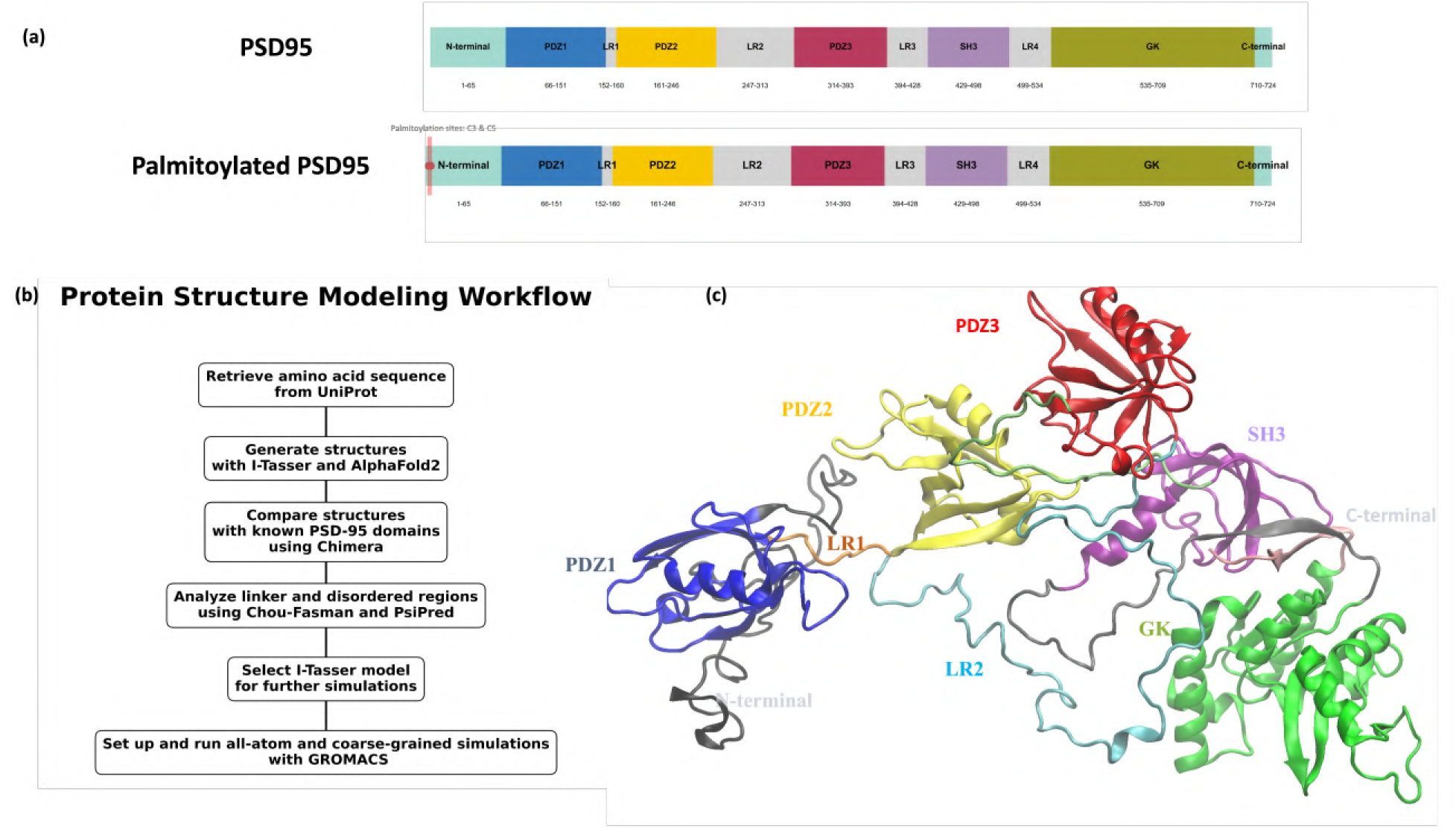
Structural organization and modeling of PSD95. (a) Domain organization of PSD95. This 724-amino acid protein comprises five domains: PDZ1 (Blue), PDZ2 (Yellow), PDZ3 (Red), SH3 (Purple), and Guanylate Kinase (Green). The N-terminal and C-termini are colored in Teal. Palmitoylation occurs at Cysteine residues 3 and 5 on the N-terminus, indicated by Red lines. (b) Protein structure modeling workflow used to generate the full-length PSD95 model. The full-length multidomain initial structure was developed using I-TASSER and subsequently used in molecular dynamics simulations. (c) 3D-structural representation of the full-length PSD95. PDZ1 (Blue) and PDZ2 (Yellow) domains connected by a short peptide linker LR1 (Orange), while the PDZ3–SH3–GK domains (Red, Purple and Green, respectively) includes more complex structural features conected by their respective linker regions. The n-terminal is colored Black and c-terminal is shown in Pink in the 3D structural representation.

## 2 Material and Methods

All input files needed to initiate molecular simulations and full trajectory data of all simulations for all systems considered in this work are publicly available on Zenodo repository. The repository can be accessed via the Zenodolink: (https://zenodo.org/uploads/15691830)

### 2.1 All-atom Molecular Dynamics Simulations Setup

Table 1 below provides the full list of all-atom systems considered in this work. Using the CHARMM-GUI interface, we constructed a total of seven systems to explore different molecular scenarios of PSD95 in our simulations. The molecular dynamics (MD) simulations were conducted using the GROMACS software package, with all-atom simulations carried out under the CHARMM36m force field.

**Table 1:**
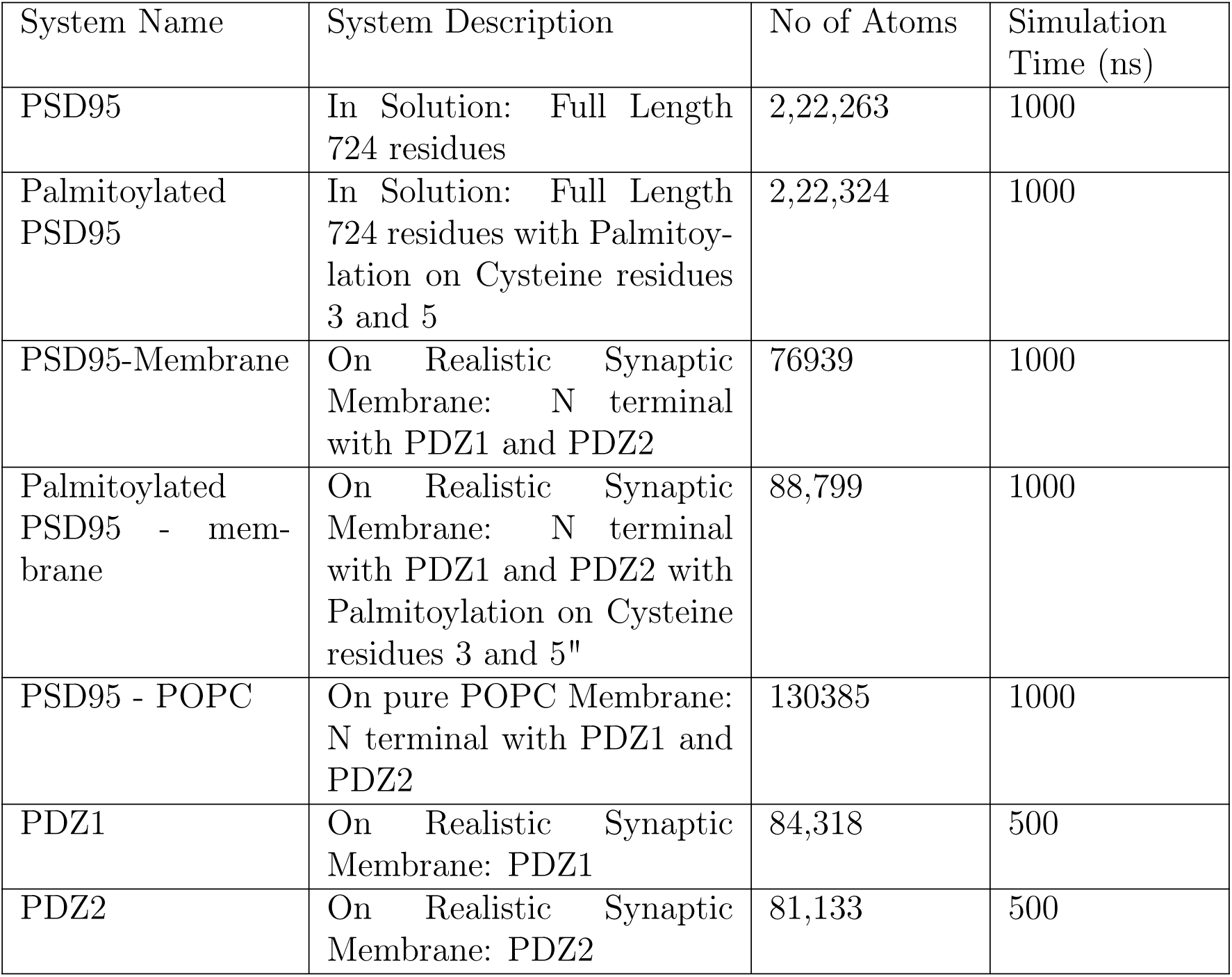
List of Simulated all atom systems.

To specifically study the impact of palmitoylation, we designed two solution-state systems. One system represented the full-length PSD95 without any modifications, while the other included PSD95 with palmitoylation. This setup was particularly useful for understanding the structural and dynamic effects of palmitoylation on PSD95 in the solution. The system was solvated in a cubic box with a minimum distance of 1 nm from the protein surface using the 3-site rigid TIP3P water model. Physiological NaCl concentration of 0.15 M was used in all simulations. Simulations were performed using Gromacs (2020.4). Energy minimization was carried out using the steepest descent algorithm for 50,000 steps to eliminate any steric clashes. The minimized structure was then sequentially equilibrated in the NVT and NPT ensembles.

Following simulations of the solution state systems, we constructed membrane-bound systems based on the lipid composition of a realistic synaptic membrane.^31–33^ The percentage composition for Phosphatidylcholine (PC), Phosphatidylethanolamine (PE), Phosphatidylserine (PS), Phosphatidylinositol (PI), Sphingomyelin (SM), Cholesterol (Chol) and Ceramide (Cer) are 35%, 30%, 12%, 3%, 2%, 17%, and 1%, respectively. These membrane systems included only the first two domains, PDZ1 and PDZ2 along with the N-terminal disordered region, since these membrane juxtaposed domains are structurally separated from the rest of PSD95 and are involved in interactions with synaptic proteins like AMPA receptors. This choice also allowed for improved computational efficiency. We created two membrane systems: one with and one without palmitoylation, to further observe the effects of this modification, but within a membrane context.

Given that our bioinformatic-based electrostatic potential analysis revealed that PDZ1 has a more negative charge than PDZ2 (suggesting that an isolated PDZ1 may be less preferred on anionic membrane than an isolated PDZ2), we aimed to see how this charge difference would influence its interaction with the membrane. We set up simulations with isolated PDZ1 and isolated PDZ2 on the synaptic membrane. As a control, we also set up a system with only neutral POPC lipids, where the unmodified PSD95 protein was introduced to study protein behavior on uncharged membrane environment.

### 2.2 Coarse-grained Molecular Dynamics Simulations Setup

Table 2 below provides the full list of coarse-grained systems considered in this work. Coarsegrained MD (CGMD) helped us explore the long-term dynamics of PSD95 on the membrane and investigate the early stages of membrane assembly features. The MARTINI 3 force field was used to represent the protein and lipid components in CG resolution. Starting from the final configurations obtained from the all-atom MD simulations, two independent systems were set up for CG simulations: one representing wild-type PSD95 and the other its palmitoylated variant. Conversion of the all-atom structures into coarse-grained representations was achieved using the Martinize2 script,^34^ which mapped the atomic coordinates into corresponding CG beads based on the MARTINI 3 force field. Subsequently, the INSANE software^35^ was employed to set up the membrane, define the simulation system boxes, solvation models, and ion concentrations.

**Table 2:**
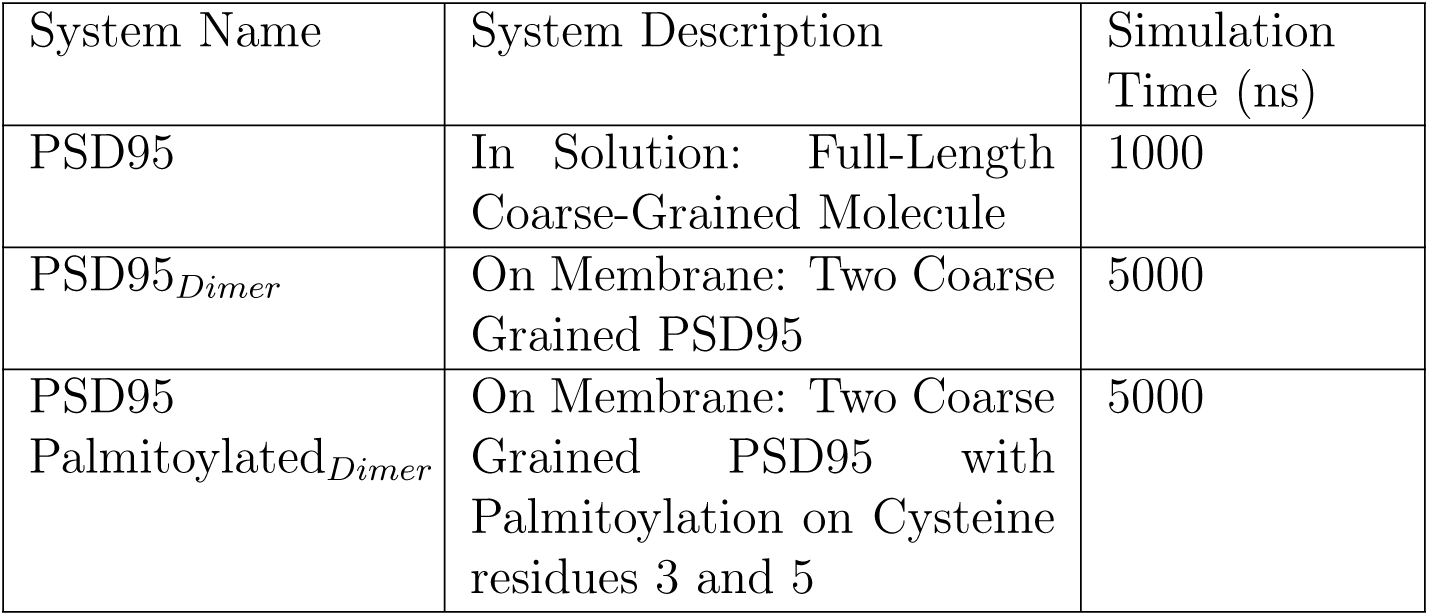
List of Simulated Coarse Grained MARTINI Systems.

| System Name | System Description | Simulation Time (ns) |
| --- | --- | --- |
| PSD95 | In Solution: Full-Length Coarse-Grained Molecule | 1000 |
| PSD95<sub>Dimer</sub> | On Membrane: Two Coarse Grained PSD95 | 5000 |
| PSD95 Palmitoylated<sub>Dimer</sub> | On Membrane: Two Coarse Grained PSD95 with Palmitoylation on Cysteine residues 3 and 5 | 5000 |

#### 2.2.1 Incorporation of palmitolytaion in Martini FF

Palmitoylation is not natively supported in the MARTINI 3 force field. To address this, we adopted the protocol introduced by Koukos et al.,^36^ which provides MARTINI 3-compatible parameters for post-translational lipid modifications. Custom mapping and topology files were integrated into our local Martinize2 installation. The mapping files (*.map) were placed in the mappings directory, and the aminoacids.ff file was extended to include parameter definitions for the palmitoyl modification.

Additionally, atomistic definitions from lipidations.rtp were incorporated into the aminoacids.rtp file within the CHARMM subfolder to enable correct modeling of the palmitoylated Cysteine residue. All atom names followed the CHARMM36m convention, and the palmitoyl moiety was covalently linked to the target Cysteine via a thioester bond. The modified protein topology was then coarse-grained using Martinize2 with the updated force field files.

#### 2.2.2 Elastic Network CG Model Optimization

To preserve the structural integrity of the protein in CG resolution, we evaluated simulations with and without an elastic network model. The radius of gyration (Rg) of both wild-type and palmitoylated systems was compared against the reference all-atom simulations. We find that inclusion of the elastic network resulted in better maintenance of native-like compactness and tertiary structure. Consequently, all subsequent CGMD simulations were performed with the elastic network enabled using standard MARTINI 3 parameters (Fig. S1).

#### 2.2.3 PSD-Bilayer setup using INSANE

Following the construction of the coarse-grained representation model for the PSD95, the protein was placed on the lipid bilayer using the INSANE (INSert membrANE) script.^35^ The bilayer was symmetric and composed of POPC:POPE:CHOL:POPS in a 40:30:20:10 ratio in both leaflets. The protein was oriented such that the palmitoyl-modified Cysteines were positioned at the membrane interface, which allowed the palmitoyl-tail to interact with the bilayer’s hydrophobic core. The simulation box was solvated, and ions were added to replicate physiological conditions.

## 3 Results and Discussion

Our results provide a comprehensive analysis of the structural and functional effects of palmitoylation on PSD95. First, we analyze the conformational changes in PSD95 and its palmitoylated form and highlight differences in overall shape and flexibility. Following this, we examine the impact of palmitoylation on the PDZ domains, particularly in relation to their stability and interaction with the membrane. Our findings indicate that palmitoylation induces a more rigid, extended structure by reducing the dynamic flipping behavior of the PDZ domains. The lipid composition of the membrane is also shown to play a crucial role in modulating these interactions, influencing both the stability and orientation of the protein. Furthermore, we explore the role of palmitoylation in facilitating the dimerization and clustering of PSD95, a key process for organizing synaptic signaling complexes. Taken together, these results underscore the importance of palmitoylation in regulating PSD95 structural state, membrane association, and functional capacity at the postsynaptic membrane.

### 3.1 Reconstructing full protein structure using integrative modeling

The process of reconstructing the full protein structure of PSD95 involved a rigorous and multi-step approach to ensure accuracy and reliability. The amino acid sequence for PSD95 was retrieved from the UniProt database (accession ID: P78352). Given the complexity of PSD95, which consists of multiple domains and disordered regions, we employed two independent computational modeling algorithms: Iterative Threading ASSEmbly Refinement (I-TASSER) and AlphaFold2. Both methods are highly regarded for their ability to predict protein structures with varying levels of precision, but they differ in their underlying methodologies.

I-TASSER, which we selected due to its robust structure prediction capabilities, combines multiple strategies to generate accurate models. The process begins with threading, where the amino acid sequence of PSD95 is aligned with known protein structures in the Protein Data Bank (PDB) to identify similar folds. It then assembles small fragments from the identified templates and refines the structure through iterative simulations to optimize the 3D conformation, accounting for physical forces that govern protein folding. This multi-step approach ensures that the final model is as accurate as possible. In parallel, we employed AlphaFold2, a state-of-the-art deep learning-based model which uses a neural network trained on vast amounts of protein sequence and structure data to predict protein fold. Unlike traditional methods that rely on template-based alignment, AlphaFold2 predicts the most likely 3D conformation based solely on the amino acid sequence, leveraging evolutionary and physical principles encoded within the network. Its novel architecture incorporates multi-sequence alignment, attention mechanisms, and complex loss functions to model the complex spatial arrangements of amino acid residues in proteins, making it particularly adept at predicting structures for proteins with no known homologous templates.

Both I-TASSER and AlphaFold2 performed well in terms of accurately predicting the individual domains of PSD95, with their structures aligning closely with known domain configurations (Fig. S2). However, discrepancies were observed in the prediction of the disordered linker regions, where the models differed in their representation of the flexible regions connecting the structured domains (Table S1). To address the discrepancies in the linker regions and further validate the structural predictions, we employed several bioinformatic methods. We employed three bioinformatic tools to validate the predicted protein structures. Chou-Fasman analysis was used to predict secondary structure propensities by analyzing the amino acid sequence, identifying regions likely to form *α* helices or *β* sheets (Fig. S3). PsiPred, a neural network-based tool, combined sequence alignments with experimentally solved structures to predict the probability of residues forming structured elements or disordered regions, providing further validation for both the I-TASSER and AlphaFold2 models (Fig. S4). To evaluate the disordered regions more thoroughly, we used PONDR, which specifically predicts intrinsically disordered segments. This allowed us to assess how well the models captured the inherent flexibility of PSD95’s linker regions. Additionally, we reviewed the existing literature on PSD95, focusing on both computational models and experimental studies that describe the interactions between its domains. This literature review provided us with context for understanding the functional relationships between the structured domains and disordered regions of PSD95.

By combining the insights from these analyses with the extensive literature review, we reached the conclusion that the I-TASSER model provided the most accurate representation of PSD95. It was particularly superior in terms of predicting the disordered linker regions and accurately reflecting the interactions between the protein’s domains. AlphaFold2 is optimized to predict stable, well-defined folded structures, relying on evolutionary data and sequence-structure relationships that work well for structured domains. However, disordered regions, which lack a fixed conformation and instead exist as dynamic ensembles, do not align with AlphaFold2’s design. AlphaFold2’s training predominantly focuses on structured proteins, and its energy minimization process biases it towards stable folds, making it less effective at modeling the flexible nature of multidomain proteins with disordered linker regions. In contrast, I-TASSER’s iterative, fragment-based approach allows for better accommodation of this flexibility, making it more adept at capturing the disorder-specific characteristics of PSD95’s linker regions.^37^ This approach of combining multiple prediction tools, structural alignment with experimental data, and literature-based validation confirmed the reliability of our model for use in simulations.

### 3.2 PSD95 and Palmitoylated PSD95 Exhibit Distinct Conformational States

Our results with all-atom molecular dynamics of PSD95 in water show that solution state PSD95 adopts distinct conformational states based on its palmitoylation status (Fig. 2(a1),(a2)). The analyses of the radius of gyration (R*_g_*) (Fig. 2(b)) demonstrates that unmodified PSD95 predominantly exists in a more compact structure. The sharp peak for the PSD95 system at lower values of R*_g_* indicates a more compact spatial arrangement of its domains. However, palmitoylation induces a significant conformational shift in PSD95, as shown by the broader and more distributed peak. The right shift suggests a larger R*_g_*, which in turn suggests a more open and extended structure for palmitoylated PSD95. The end-to-end distance analysis (Fig. 2(c)) further supports these results. Unmodified PSD95 shows a concentrated peak at a shorter distance, which is consistent with the compact state observed in the radius of gyration. In contrast, palmitoylated PSD95 displays a broader range of end-to-end distances, reflecting a more dynamic and extended structure. The movie of the trajectories for PSD95 and pal-PSD95 are available as movieS1a and movieS1b, respectively in the SI PPTX file included in the zenodo repository (https://doi.org/10.5281/zenodo.15691830).

**Figure 2:**
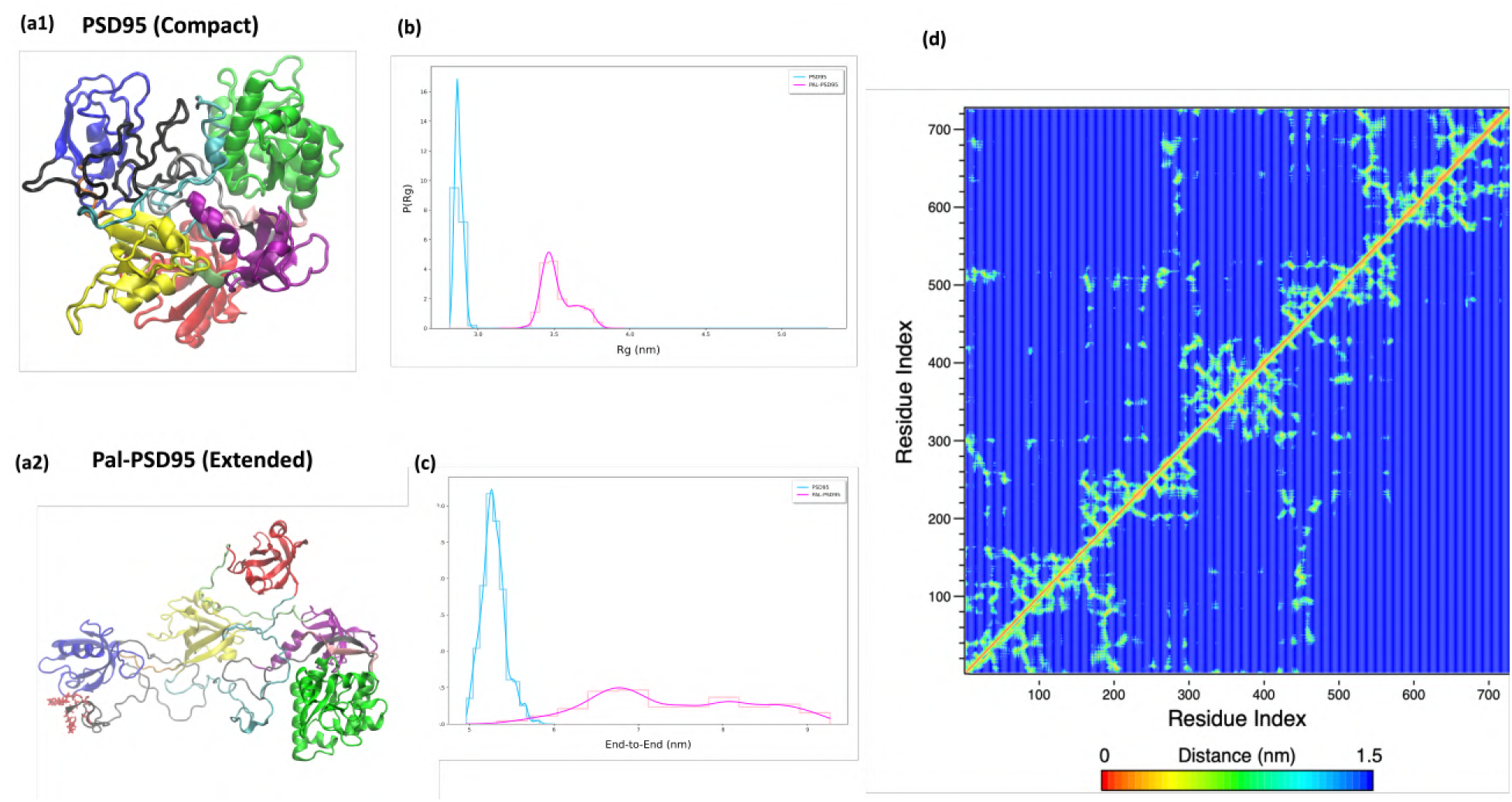
Structural properties of PSD95 and Pal-PSD95 in solution. (a1) Structural model of PSD95 in its compact conformation, showing tight packing of domains. (a2) Structural model of Pal-PSD95 in its extended conformation following palmitoylation. (b) Radius of gyration indicating overall compactness of unmodified PSD95 (blue) and palmitoylated PSD95 (pink). (c) End-to-end distance measuring the spatial separation between the N- and C-termini of PSD95 (blue) and Pal-PSD95 (pink). (d) Residue–residue contact maps of PSD95 (left upper diagonal) and pal-PSD95 (right lower diagonal), showing differences in intramolecular contacts and folding patterns induced by palmitoylation.

Finally, the contact maps (Fig. 2(d)) highlight the impact of palmitoylation on the residue-residue interactions within PSD95. The contact map of the unmodified PSD95 (left upper diagonal) reveals a tighter clustering with several off-diagonal contacts having noticeable frequencies of contacts. On the other hand, palmitoylated PSD95 (right lower diagonal) shows a more diagonal centric dispersion pattern of contacts, aligning with the idea of an extended conformation. The redistribution of interactions suggests that palmitoylation not only modifies the overall shape of PSD95 but also alters its internal domain interactions (Fig. S5).

Collectively, these results indicate that palmitoylation induces a significant structural shift in PSD95 in the solution state and transforms it from a compact to an extended state. This structural rearrangement is likely crucial for PSD95’s function at the postsynaptic membrane, potentially enhancing its role in scaffolding and interacting with other synaptic components. The flexibility conferred by palmitoylation may therefore be a key regulatory mechanism to modulate synaptic stability and plasticity.

### 3.3 Palmitoylation Induces PDZ Domain Flipping on the membrane

Our all-atom molecular dynamics PSD95 simulations with membrane bilayer reveal that palmitoylation of PSD95 leads to domain rearrangement. As seen in (Fig. 3(a1)), the unmodified PSD95 shows dynamic movement between the PDZ1 (Blue) and PDZ2 (Yellow) domains. Specifically, during the simulation of unmodified PSD95, the PDZ1 domain tends to move away from the membrane, while the PDZ2 domain shifts closer. This “flipping” behavior indicates a level of flexibility in the unmodified state, where the domains are more mobile and unstable. However, once PSD95 undergoes palmitoylation, the dynamics of the PDZ domains change dramatically. Palmitoylated PSD95 remains in a more extended state throughout the simulation, with both PDZ1 (Blue) and PDZ2 (Yellow) showing less movement and more stable positioning (Fig. 3(a2)). This reduction in domain flipping upon palmitoylation suggests that the addition of palmitoyl groups stabilizes the overall structure of PSD95, anchoring the protein more tightly to the membrane and reducing the flexibility of the PDZ domains.

**Figure 3:**
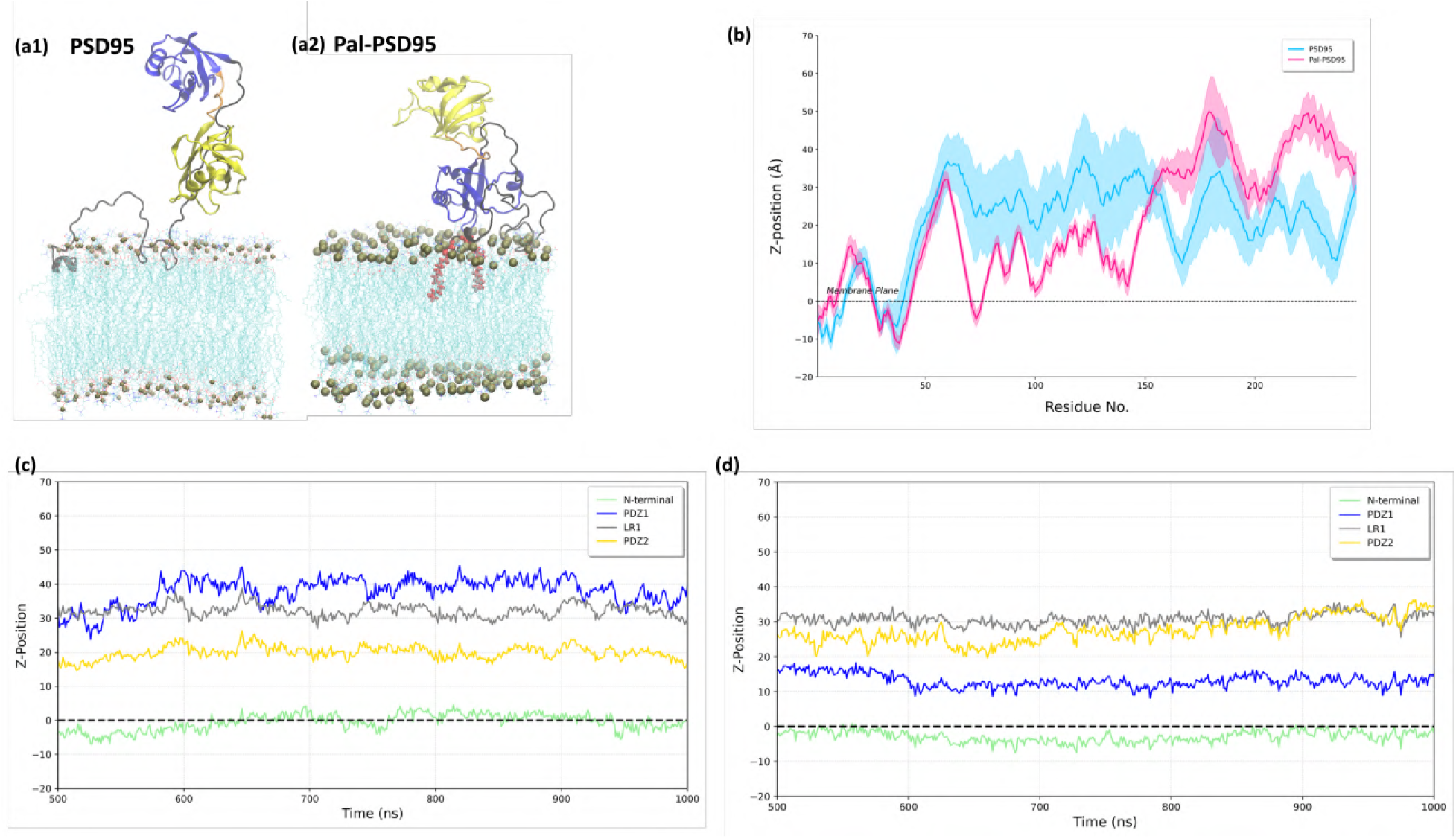
Membrane interaction dynamics of PSD95 and Pal-PSD95. (a1) Unmodified PSD95 on the membrane shows dynamic behavior between PDZ1 (Blue) and PDZ2 (Yellow) domains. During the simulation, PDZ1 tends to move away from the membrane while PDZ2 shifts closer, resulting in domain “flipping.” (a2) Palmitoylated PSD95 adopts and maintains an extended conformation, with both PDZ1 and PDZ2 stably anchored to the membrane throughout the simulation. (b) Ensemble-averaged membrane distance profiles for each residue reveal reduced variability and more consistent membrane association in the palmitoylated system (pink) compared to unmodified PSD95 (Blue). Time-dependent domain wise membrane distance plots for (c) PSD95 and (d) pal-PSD95 show sustained proximity of PDZ1 to the membrane in pal-PSD95 system while PDZ1 is distal and PDZ2 is membrane proximal in unmodified PSD95.

In the ensemble data shown in Fig. 3(b), we observe a clear difference between unmodified PSD95 and palmitoylated PSD95 with respect to their residue-wise membrane association. The shaded areas represent standard deviations around the average position, capturing variations in domain movement over time. This figure further highlights that palmitoylation strengthens PSD95’s association with the membrane and alters the domain-flipping behavior seen in the unmodified protein. Fig. 3(c,d) show time-dependent membrane distance plots that trace the positional evolution of different domains throughout the simulations of two systems. Additionally, we find increased lipid contacts for the N-terminal and PDZ1 domain of the palmitoylated PSD95, with no membrane contact observed for the PDZ2 domain (Fig. S6). Overall, our data suggest that the stabilized, extended configuration of the palmitoylated form likely enhances PSD95’s scaffolding role at the synapse by keeping the protein in a more rigid, functionally optimized state, thereby minimizing the domain dynamics and flexibility observed in the unmodified version. The movie of the trajectories for membrane bound PSD95 and pal-PSD95 are available as movieS2a and movieS2b, respectively in the SI PPTX file included in the zenodo repository (https://doi.org/10.5281/zenodo.15691830).

This was quite an interesting and unexpected observation since the PDZ1 domain is positioned between membrane proximal disordered N-terminal region and PDZ2 domain yet PDZ2 prefers to stay closer to the membrane rather than PDZ1. To further investigate why this behavior occurs, we placed the individual isolated PDZ1 and PDZ2 domains on the membrane and ran separate all-atom molecular dynamics simulations for 500 nanoseconds each (Fig. 4 (a1,a2). Our results indicate differences in how each domain interacts with the membrane: On its own, PDZ1 is less favored on the membrane than PDZ2. The z-plot for isolated PDZ1 and PDZ2 are shown in Fig. 4(b) and Fig. 4(c), respectively. The domain flipping can be explained by the electrostatic properties of each domain in relation to the membrane. Because the membrane carries a negative charge, it doesn’t tolerate PDZ1 domain as it is interspersed with negative residues resulting in its tendency to move away from the membrane. In contrast, PDZ2 has a more neutral charge distribution, which is less affected by the membrane’s negative charge that allows it to remain stably anchored to the membrane. In the unmodified PSD95, the 65-amino acid long disordered N-terminal region provides enough steric freedoma for the distal PDZ2 to approach the membrane. When palmitoylated, the N-terminal disordered region is sterically restricted and this allows the adjacent PDZ1 to be closer to the membrane. These findings indicate that there is a balance between the charge complementarity and anchoring by modification that play a significant role in the domain-specific membrane-binding flipping behavior of PSD95 in absence and presence of the palmitoylation modification. The movie of the trajectories for isolated PDZ1 and PDZ2 are available as movieS3a and movieS3b, respectively in the SI PPTX file included in the zenodo repository (https://doi.org/10.5281/zenodo.15691830).

**Figure 4:**
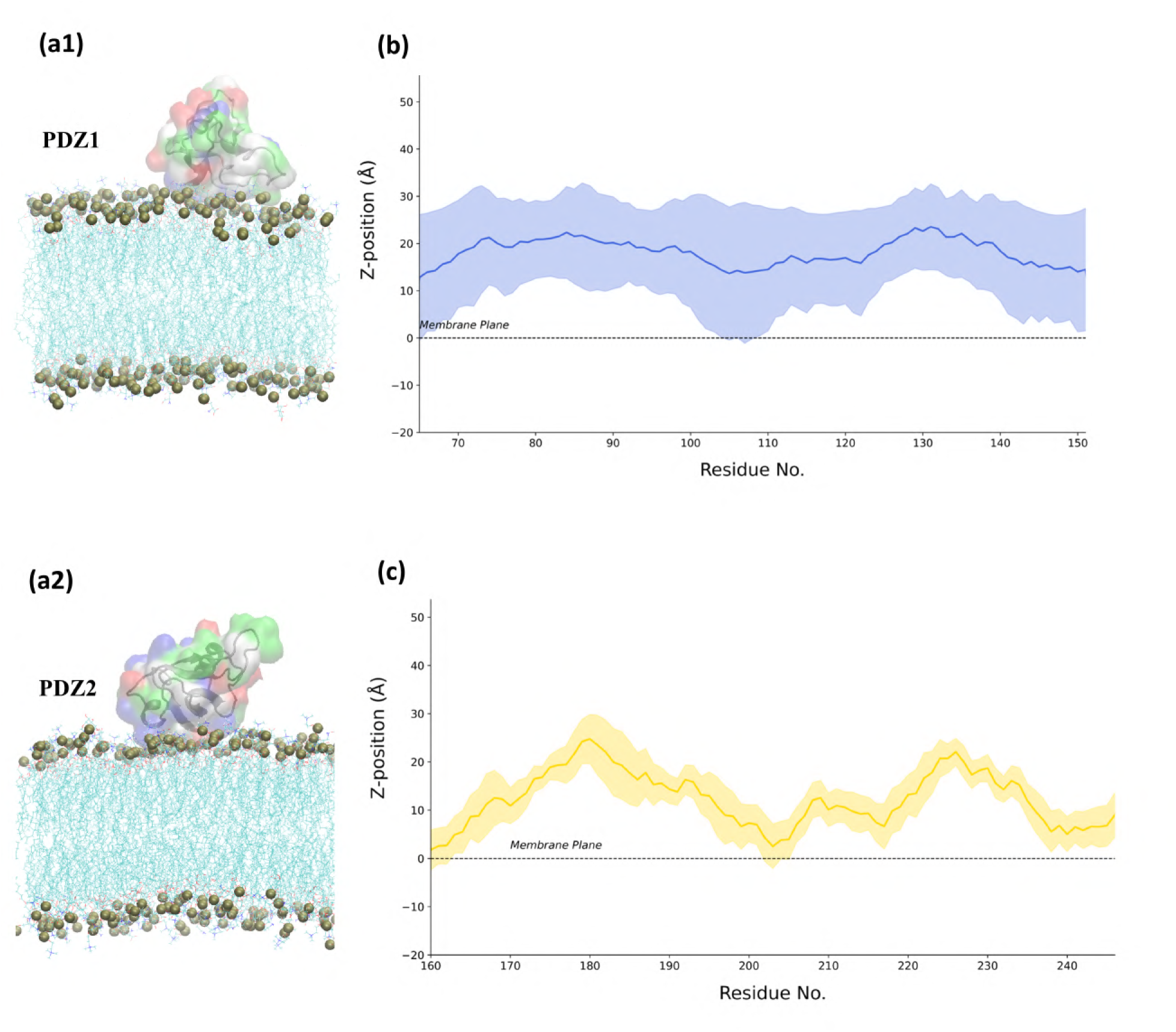
Membrane interaction profiles of isolated PDZ1 and PDZ2 domains. (a1, b) Simulation snapshots showing the spatial orientation of isolated PDZ1 and the corresponding residue-wise membrane distance profiles relative to the membrane. Isolated PDZ1 exhibits minimal and unstable membrane association, likely due to its higher surface negativity. (a2, c) Simulation snapshots showing the spatial orientation of isolated PDZ2 and the corresponding residue-wise membrane distance profiles relative to the membrane. PDZ2 displays stronger and more stable membrane interactions, indicating more favorable electrostatic compatibility. These differences help explain the domain-flipping behavior observed in full-length PSD-95 and highlight the contribution of intrinsic domain properties to membrane binding.

### 3.4 Lipid Composition Affects PSD95 Interaction with Membranes

In order to establish the interplay between electrostatic interactions and anchoring by palmitoylation, we carried out simulations with different lipid compositions. The lipid composition of the membrane significantly influences how PSD95 interacts with it, affecting both the stability and orientation of the protein. Negatively charged lipids, which are commonly found in synaptic membranes, can repel similarly charged regions of PSD95, impacting the stability of interactions.

This behavior was further clarified through simulations with a POPC-only membrane, which carries a more neutral charge. In the POPC membrane environment, both PDZ1 and PDZ2 showed similar stability, suggesting that charge interactions within synaptic membranes, rather than lipid composition alone, contribute to the distinct flipping behavior observed in PSD95 domains (Fig. 5). Also, he movie of the trajectories for PSD95 on POPC membrane is available as movieS4 in the SI PPTX file included in the zenodo repository

**Figure 5:**
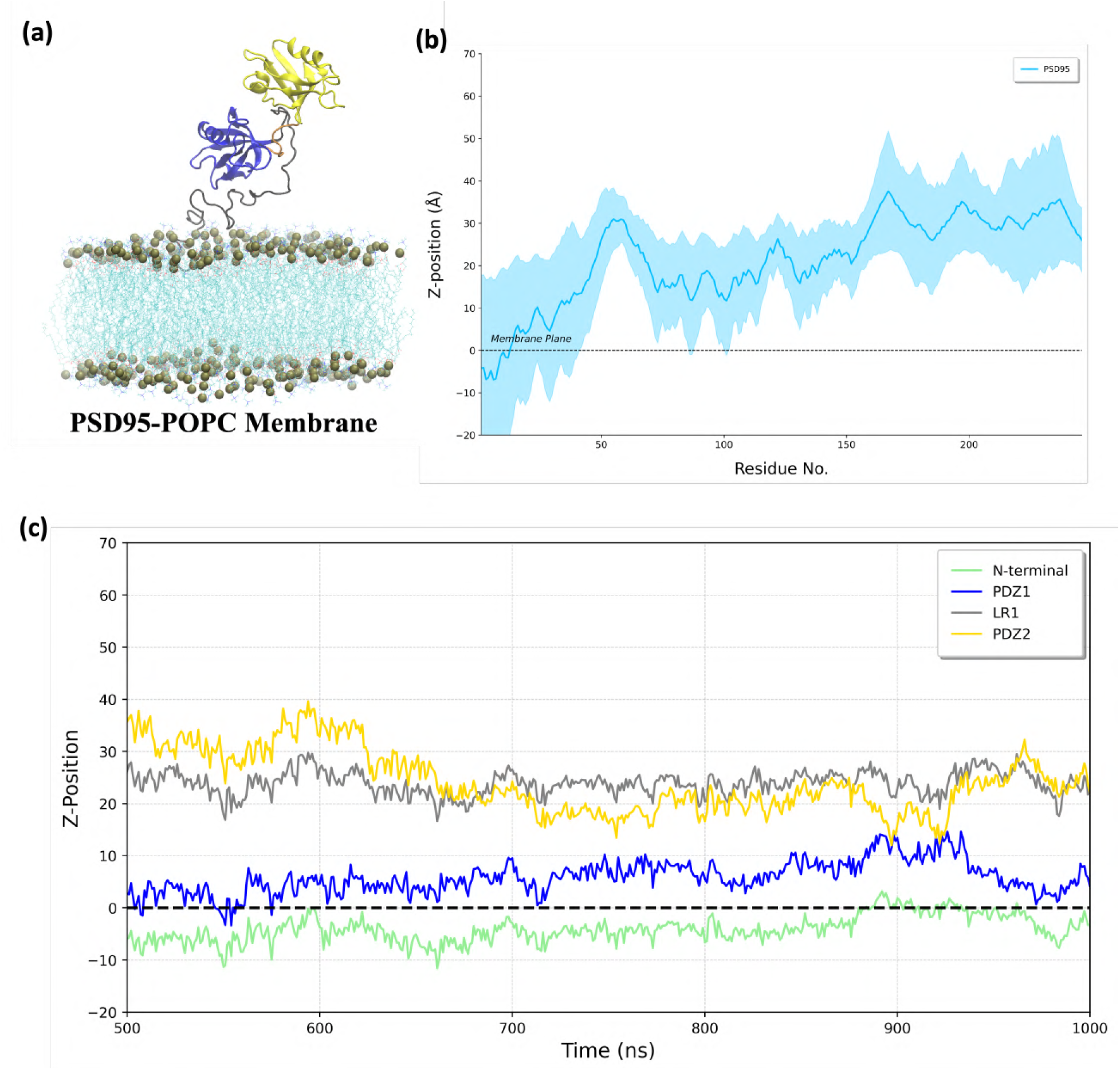
Membrane composition influences PSD95 domain orientation. (a) Simulation snapshot of PSD95 on a POPC-only membrane shows minimal domain reorientation, lacking the distinct domain flipping observed in synaptic membrane environments. (b) Ensemble-averaged membrane distance profile shows decreased domain flipping compared to synaptic membranes. (c) Time-dependent membrane distance of individual domains (N-terminal, PDZ1, linker, and PDZ2) highlights the reduced flipping behavior in the POPC membrane.

### 3.5 Palmitoylation Promotes the Formation of Robust PSD95 Dimers on the Membrane

To further investigate whether palmitoylation facilitates PSD95 cluster formation at the membrane, we carried out coarse-grained MARTINI simulations of two PSD95 molecules on a synaptic lipid membrane. Coarse-grained simulations allowed us to access longer timescales and assess the stability of dimerization events beyond the limits of all-atom simulations.

In the palmitoylated system, the distance between the two proteins rapidly decreased and remained stable throughout the simulation (Fig. 6(a1)), indicating the formation of a stable dimer. The contact map generated at 5 µs shows extensive and sustained inter-protein interactions (Fig. 6(b,c), particularly involving residues from the PDZ1 and PDZ2 domains. These results suggest that palmitoylation promotes not only membrane anchoring but also a favorable orientation for lateral assembly and inter-molecular contacts. In contrast, simulations of unpalmitoylated PSD95 molecules under identical conditions exhibited fluctuating and unstable distances between the two proteins (Fig. 6(a2)), and the corresponding contact map revealed sparse or transient inter-protein interactions (Fig. 6(d,e)). Without palmitoylation, PSD95 molecules appeared to interact transiently before drifting apart, failing to form a stable dimeric arrangement. The movie of the trajectories for PSD95 and pal-PSD95 dimers are available as movieS6a and movieS6b, respectively in the SI PPTX file included in the zenodo repository (https://doi.org/10.5281/zenodo.15691830). This indicates that palmitoylation is a key requirement for sustained membrane-bound self-association. All simulations were performed using a synaptic lipid composition that includes negatively charged lipids. Taken together, these data demonstrate that palmitoylation serves as a minimal structural requirement for PSD95 clustering, and likely acts as a molecular switch that enables both anchoring and lateral interactions on the membrane. These findings support the hypothesis that palmitoylation directly contributes to the early stages of postsynaptic scaffold formation.

**Figure 6:**
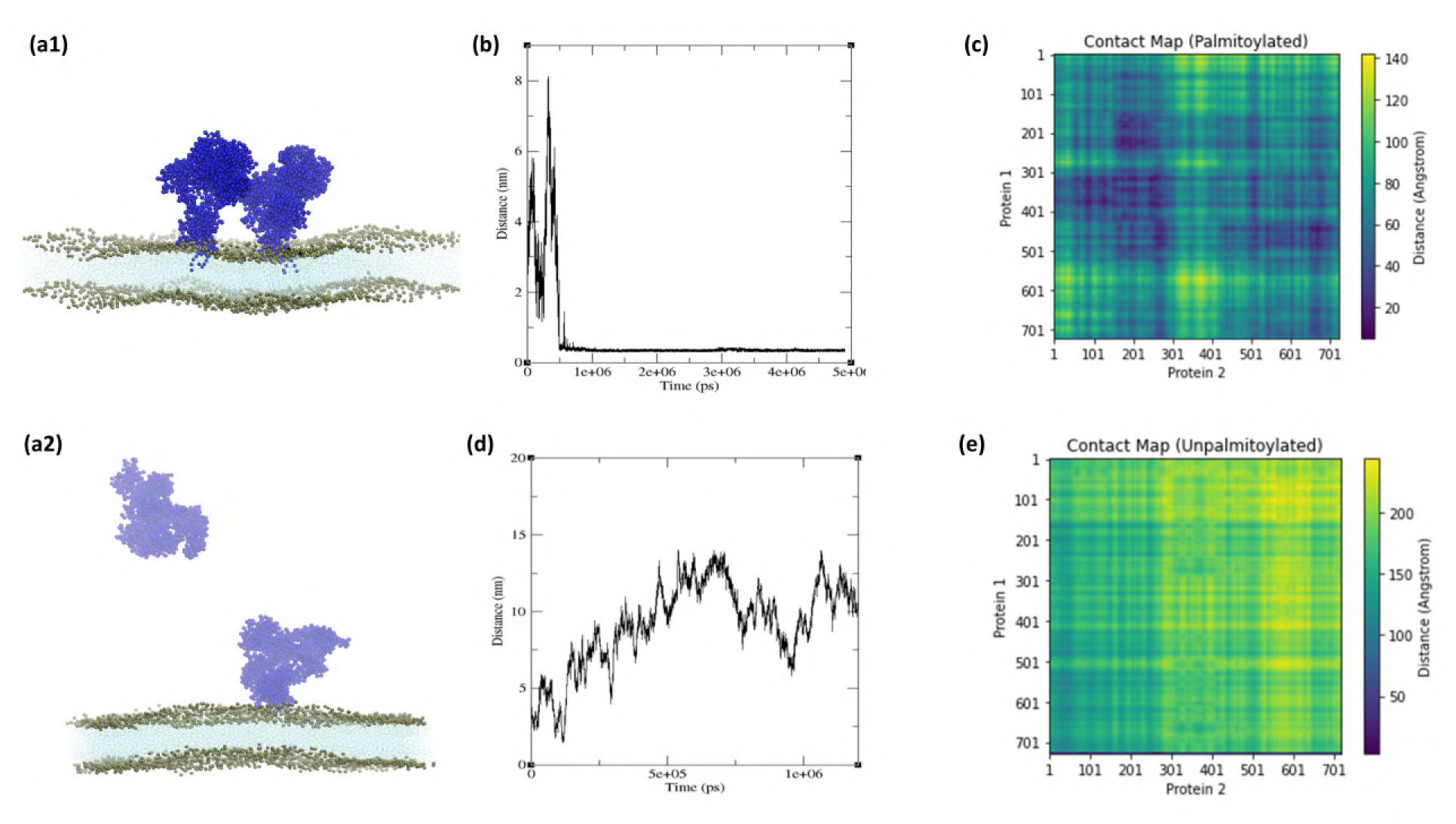
Palmitoylation promotes membrane-anchored dimerization of PSD95 in coarse-grained simulations. (a1–c) Palmitoylated PSD95: (a1) Snapshot of the Martini simulation at the end of a 5-microsecond run, showing two palmitoylated PSD95 molecules (blue) stably associated on the membrane. (b) Distance between the centers of mass of the two palmitoylated PSD95 molecules over time, showing stable proximity and dimer formation. (c) Contact map for the palmitoylated dimer at 5 µs, indicating sustained inter-protein interactions. (a2, d–e) Unpalmitoylated PSD95: (a2) Snapshot of the Martini simulation at the end of a 1-microsecond run. (d) Distance plot for unpalmitoylated PSD95 dimers shows unstable, transient interaction and eventual separation. (e) Contact map for the unpalmitoylated system reveals minimal or no persistent interactions between proteins. The movie files of the two trajectories are available in the PPTX file included in the zenodo repository

## 4 Conclusions

Our results reveal how palmitoylation influences the conformation, membrane association, and clustering of PSD95, providing key insights into how this post-translational modification modulates synaptic protein dynamics and impacts synaptic structure and function.^38^ We observed a significant conformational change induced by palmitoylation, which promotes an extended configuration in PSD95. This extended conformation likely enhances its ability to interact with the postsynaptic membrane and other synaptic proteins, increasing the surface area available for interactions.^13,39–41^ These conformational changes can facilitate the organization of key neurotransmitter receptors, such as NMDA and AMPA receptors, and thus improve synaptic signaling efficiency.^42–44^ Palmitoylation, therefore, appears to play a crucial role in optimizing the scaffolding functions of PSD95, contributing to effective synapse organization.^45–47^

In addition to conformational changes, our simulations revealed that palmitoylation strengthens PSD95’s membrane association and stabilizes its position, particularly with respect to the PDZ2 domain, which remains closer to the membrane when palmitoylated.**^?^** This proximity may be important for synaptic protein interactions, allowing the PDZ domains to efficiently bind transmembrane proteins like neurotransmitter receptors. The palmitoyl groups seem to act as membrane anchors, enhancing PSD95’s lateral mobility and stabilizing its extended state.^48^ This stabilization is essential for synaptic function as it ensures proper protein positioning for effective signal transduction. Our findings suggest that palmitoylation is not just a structural modification but a functional one that fine-tunes PSD95’s ability to organize and regulate synaptic proteins.^45^

The coarse-grained simulations further demonstrated that palmitoylation aids in robust clustering and dimerization of PSD95 on the membrane.^49^ This clustering is critical, as PSD95 clusters act as scaffolds for other postsynaptic proteins, helping to organize synaptic signaling complexes. By promoting stable dimerization and subsequent cluster formation, palmitoylation enhances the structural integrity of synapses, making them more capable of supporting efficient neurotransmission.^50^ The observed clustering behavior suggests that palmitoylation is a key regulatory mechanism that governs how PSD95 assembles at synapses, which has implications for understanding synaptic plasticity and the molecular organization underlying it.

Our findings also indicate that the lipid composition of the membrane influences PSD95’s behavior. Specifically, the presence of phosphatidylserine (PS) and cholesterol in the synaptic membrane appears to enhance the clustering of palmitoylated PSD95.^51,52^ The negatively charged PS likely interacts with positively charged regions on PSD95, stabilizing its membrane association, while cholesterol may create lipid microdomains that promote protein clustering. This highlights the importance of the lipid environment in modulating PSD95’s membrane interactions and underscores how the synaptic membrane composition contributes to the overall organization of synaptic proteins.

Impaired PSD95 clustering has been linked to neurodevelopmental disorders such as schizophrenia and autism, where synaptic signaling is disrupted.^53^ Our study suggests that disruptions in the palmitoylation pathway could contribute to such synaptic deficits. Understanding the role of palmitoylation in PSD95’s behavior could pave the way for therapeutic interventions targeting synaptic dysfunction, potentially offering novel approaches for treating disorders characterized by impaired synaptic signaling.

In conclusion, this study sheds light on the molecular mechanisms by which palmitoylation regulates PSD95’s structure, membrane association, and clustering. While our findings provide significant insights into the role of palmitoylation in PSD95 function, the next logical step is to investigate how these mechanisms are influenced by interactions with stargazin, to better understand the broader assembly and stability of AMPA receptor complexes. ^54,55^

Future studies should combine MD simulations and experimental approaches to elucidate how PSD95 palmitoylation modulates its interaction with stargazin. Specifically, it will be critical to determine whether palmitoylation-induced conformational changes in PSD95 enhance its affinity for stargazin and whether this interaction further stabilizes receptor clustering and synaptic organization.

Additionally, exploring the cooperative role of stargazin in PSD95 clustering and nanodomain formation could provide a more comprehensive understanding of the molecular architecture of the postsynaptic density. Given that stargazin itself is regulated by phosphorylation, future investigations should examine the interplay between PSD95 palmitoylation and stargazin phosphorylation in modulating the dynamics of receptor anchoring and synaptic plasticity.

Finally, it would be valuable to expand these studies to more complex synaptic environments, such as those involving other TARPs, NMDA receptors, and auxiliary scaffolding proteins like GKAP or Shank.^56,57^ These multi-protein systems could reveal how PSD95’s palmitoylation integrates with stargazin binding to orchestrate the assembly of diverse signaling complexes. Such insights would deepen our understanding of synaptic regulation and its implications for synaptic disorders, potentially uncovering new therapeutic targets for various neuropsychiatric conditions.

## Acknowledgement

AS acknowledges the financial support from the Indian Institute of Science (IISc) and the high-performance computing facility “Beagle” that was set up from grants by the erstwhile IISc-DBT partnership programme. DN and AS thank the DST for the National Supercomputing Mission grants (DST/NSM/R&D-HPC-Applications/Extension Grant/2023/27). AS also acknowledges the DST FIST program that supports the department infrastructure. AS thanks the DBT-Wellcome Trust India Alliance (Grant number: IA/TSG/21/1/600245). AS also thanks the DBT National Network Project (NNP) grant (BT/PR40323/BTIS/137/78/2023) and the Matrics grants (MTR/2023/001040) from the Science and Engineering Board (SERB), India. DN also thanks the DBT-Wellcome Trust India Alliance grant (IA/S/23/2/507005) and his ANRF grant (CRG/2022/002726).

## Author contributions

DN and AS conceptualized the project. AS designed the research. PD performed the research and all the simulations. PD analyzed the data with help from DN and AS. AS supervised the study. PD prepared the first draft of the paper with inputs from AS. DN and AS polished the draft.

## 5 Conflict of interest

The authors declare that they have no potential conflict of interest.

## 6 Data availability statement

Our code, models, and curated datasets as well all the movie files of all the MD trajectories are publicly available at the following Zenodo repository https://doi.org/10.5281/zenodo.15691830.

## Supporting Information

### List of movie files

The movie files from the various MD trajectories are embedded in the PPTX file included in the zenodo repository (https://doi.org/10.5281/zenodo.15691830).

1. **movieS1a and movieS1b**: Two movie files with solution state PSD95, w w/o PAL.
2. **movieS2a and movieS2b**: Two movies with PDZ1+PDZ2 on membrane w w/o PAL). with POPC membrane.
3. **movieS3a and movieS3b**: Isolated PDZ1 and PDZ2 on membrane.
4. **movieS4**: PSD95 on POPC membrane.
5. **movieS5a and movieS5b**: CGMD custering with PSD95 dimers on the membrane - w and w/o PTM.

**Table 1:** List of linker regions and terminal segments in PSD95.

| **Region** | **Residue Range** |
| --- | --- |
| N-terminal | 1–65 |
| LR1 | 152–160 |
| LR2 | 247–313 |
| LR3 | 394–428 |
| LR4 | 499–534 |
| C-terminal | 710–724 |

### Secondary Structure Propensities of Linker Regions

PSD95 is a multi-domain protein connected by several intrinsically disordered linker regions between well-folded domains. To better understand the conformational behavior of these linkers, we analyzed their secondary structure propensities. These flexible segments play a key role in mediating domain dynamics and structural rearrangements. We used I-TASSER for initial structure prediction, as it effectively incorporates secondary structure predictions and provides conformations that are consistent with known folding patterns of ordered and disordered regions. This approach allowed us to model the overall topology of PSD95 while preserving the flexibility and structural variability of its inter-domain linkers.

**Fig. S 1:**
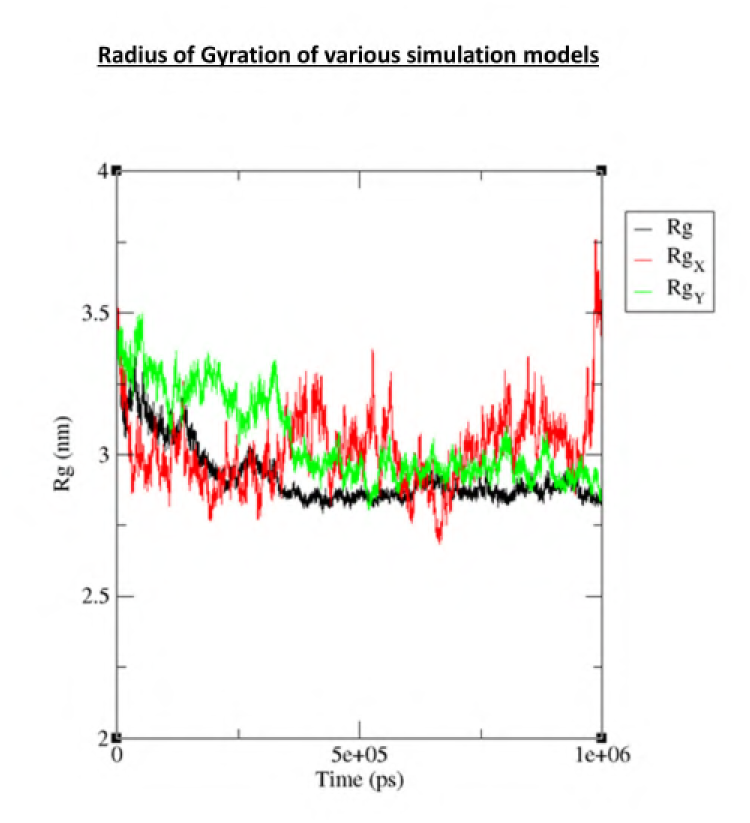
Radius of gyration (Rg) across different simulation models. The plot compares the Rg values over a 1 s trajectory for three distinct setups. The black line represents the all-atom simulation, serving as a reference for an accurate dynamics. The red line shows a MARTINI simulation without an elastic network, and the green line represents a MARTINI simulation with an elastic network.

**Fig. S 2:**
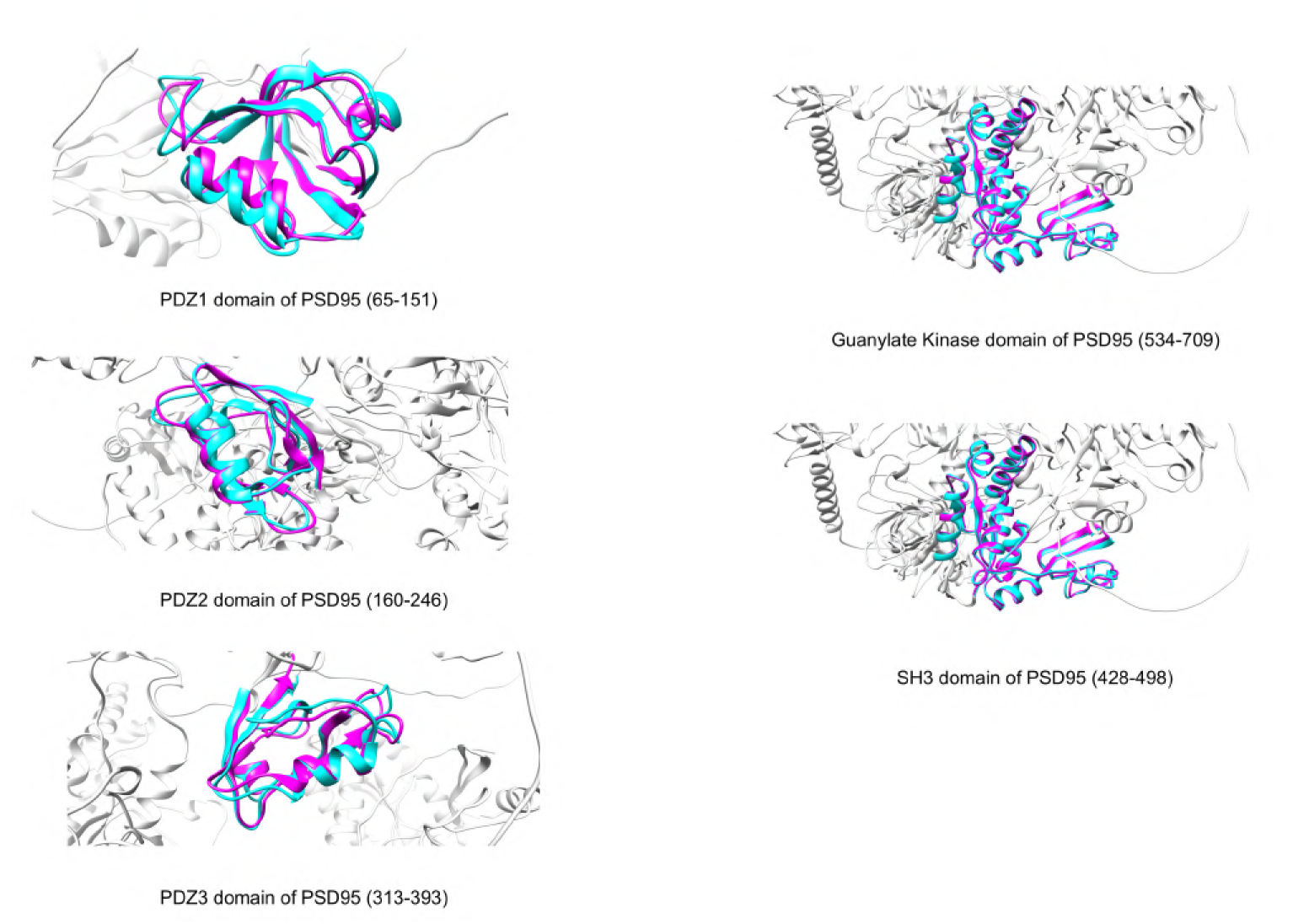
Structural alignment of predicted models with reference structures for PSD95 domains. Structural alignments of AlphaFold models (cyan) and I-TASSER models (magenta) with reference PDB structures are shown for five key domains of PSD95: PDZ1 (residues 65–151), PDZ2 (residues 160–246), PDZ3 (residues 313–393), SH3 (residues 428–498), and Guanylate Kinase (GK) domain (residues 534–709). The reference structures are depicted in gray to provide structural context within the full-length protein. The overlay of predicted models demonstrates high structural similarity across all domains.

**Fig. S 3:**
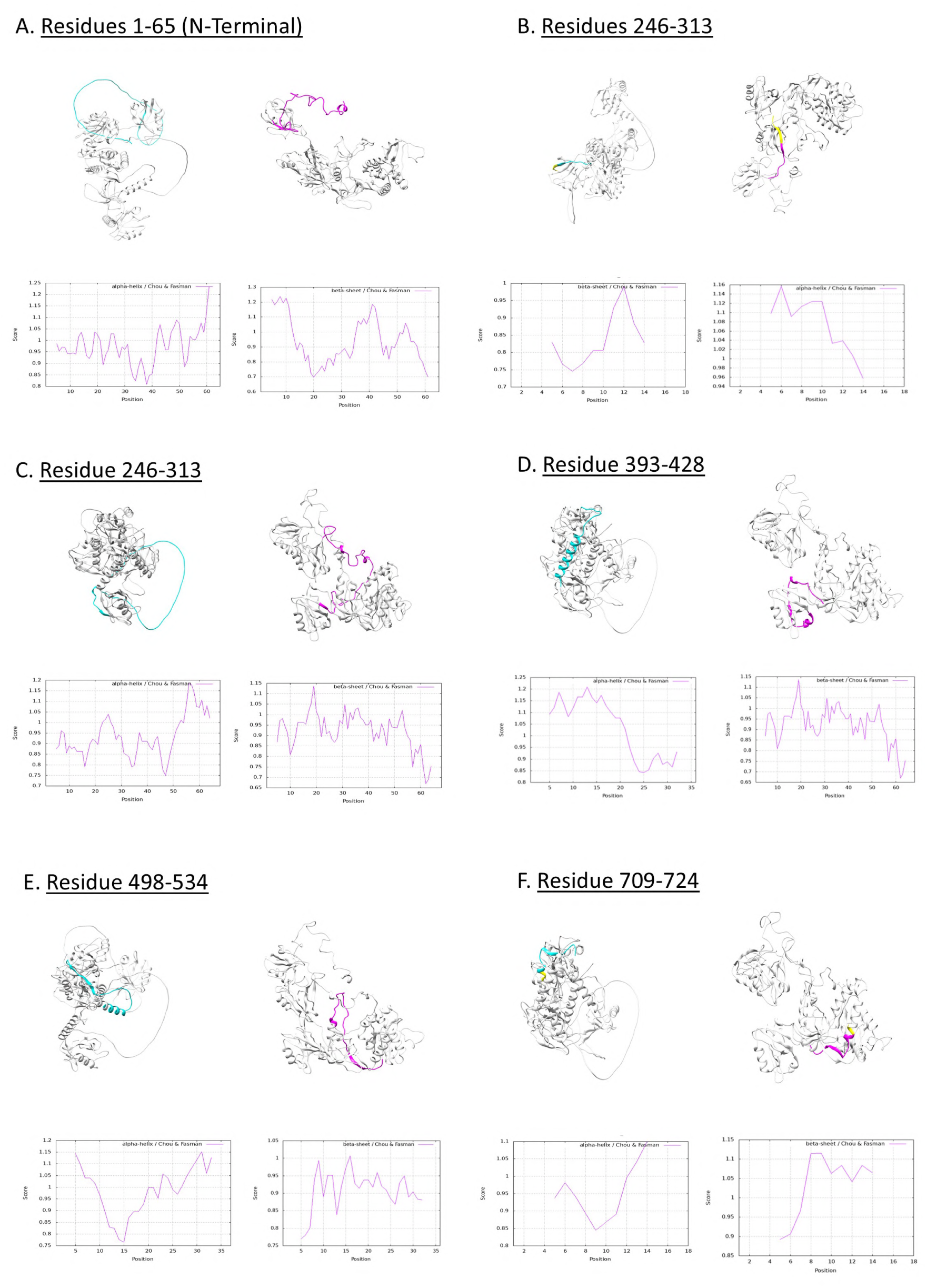
Comparison of AlphaFold and I-TASSER predicted structures for selected regions of a protein. Panels A–F represent different residue segments, with AlphaFold predictions shown in cyan and I-TASSER predictions in magenta. The corresponding Chou-Fasman secondary structure propensity plots are shown b_6_elow each structure, illustrating the likelihood of alpha-helix, beta-sheet, or coil formation across the sequence. This comparison highlights structural variations between the two prediction methods and their alignment with secondary structure propensities.

**Fig. S 4:**
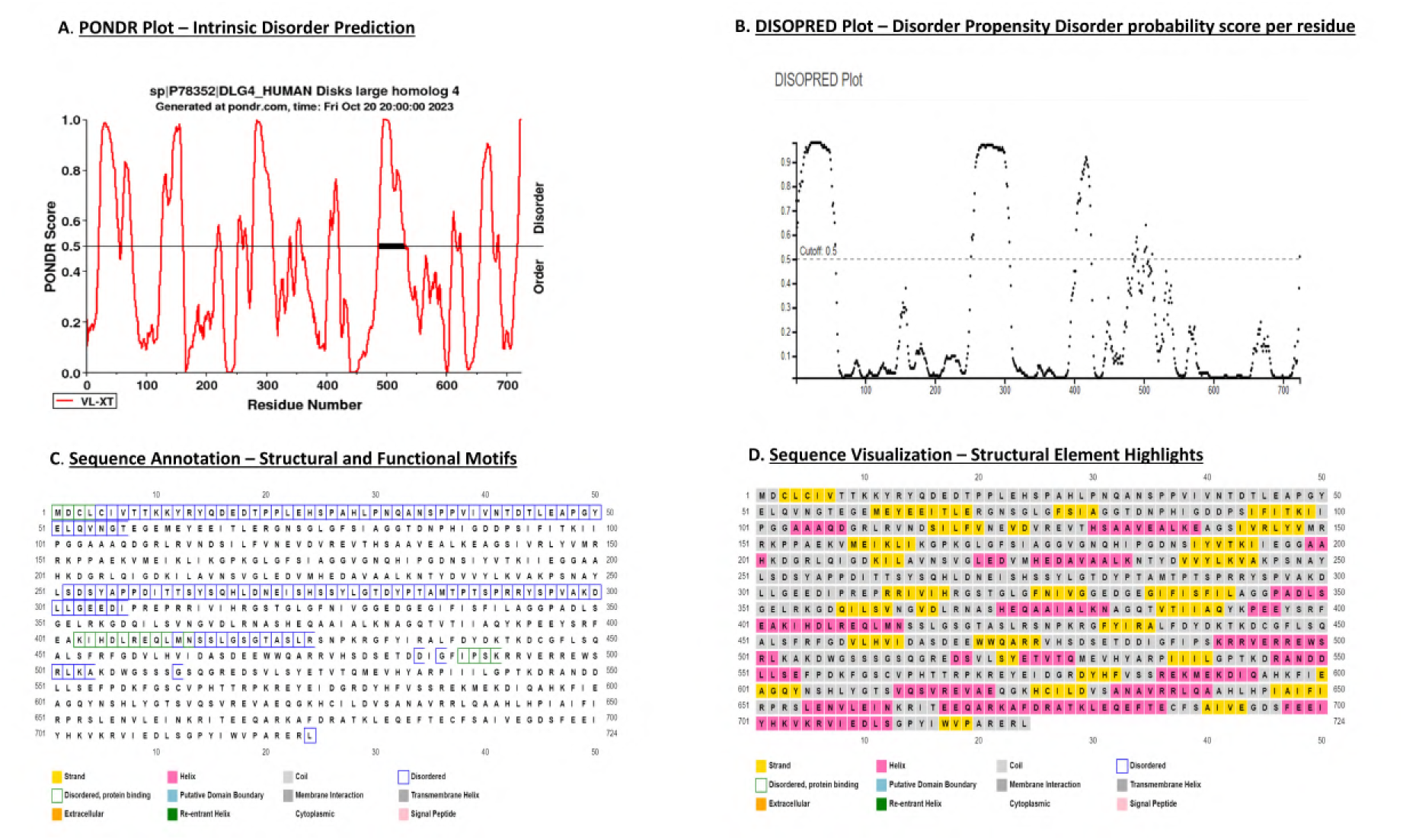
Disorder prediction and sequence-based annotation of PSD95. (A) PONDR Plot shows intrinsic disorder prediction across the PSD95 protein sequence using the VL-XT algorithm. A score above 0.5 indicates predicted disordered regions. (B) DISOPRED Plot displays disorder propensity scores per residue. Regions scoring above the 0.5 cutoff are considered disordered. (C) Sequence annotation with structural and functional motifs, including secondary structures (helix, strand, coil), domain boundaries, disordered regions, and membrane-related features, using UniProt annotations. (D) Sequence visualization highlighting structural elements such as helices (pink), strands (yellow), disordered regions (gray), and signal peptides, providing a comprehensive view of sequence-derived structural features. Color codes are indicated in the legend below for reference.

**Fig. S 5:**
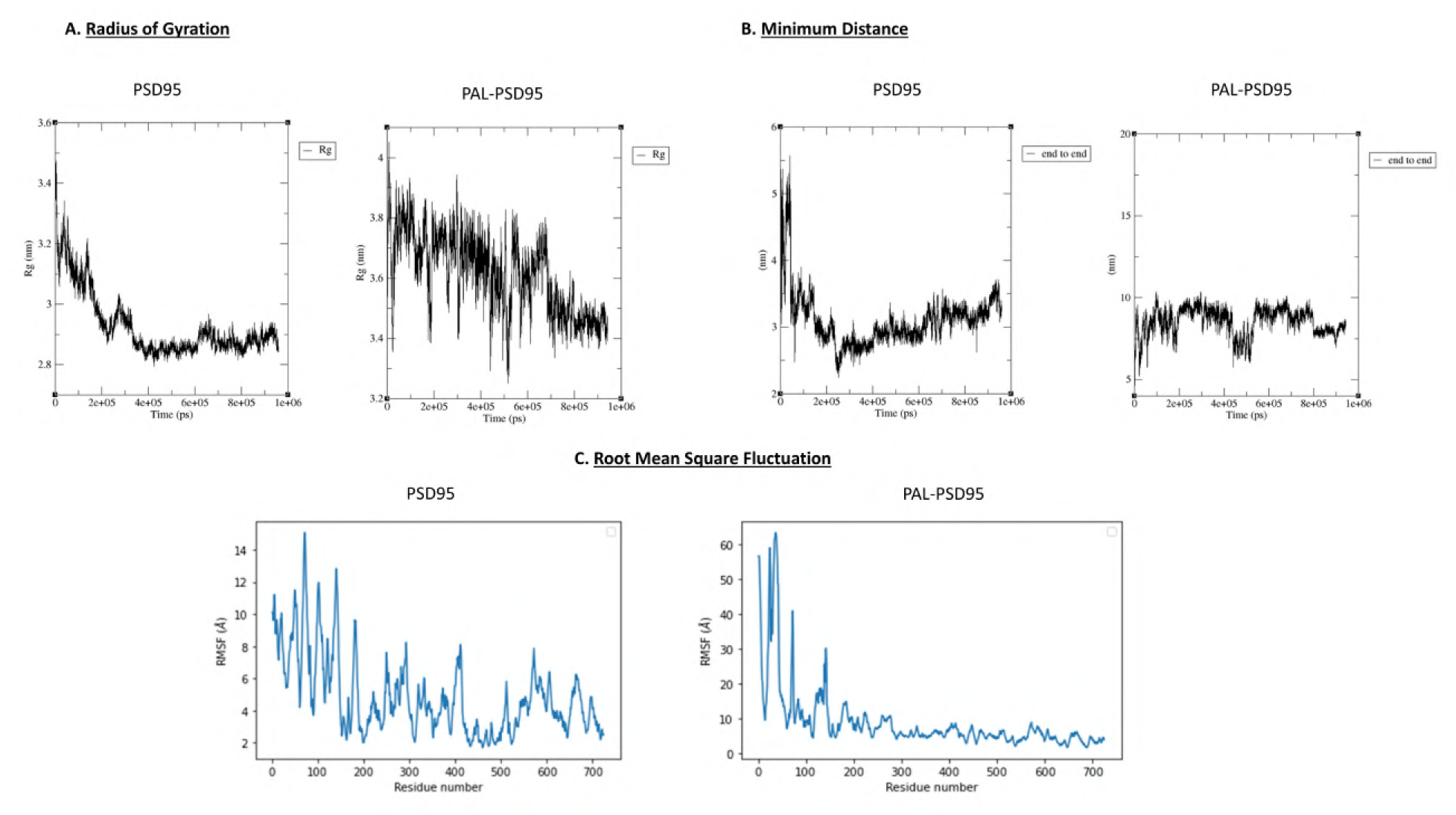
Conformational dynamics of PSD95 and palmitoylated PSD95. (A) Radius of Gyration (Rg): Time evolution of the radius of gyration (Rg) of PSD95 (left) and PAL-PSD95 (right) over a 1 s simulation. PAL-PSD95 exhibits a higher average Rg, suggesting an overall more extended conformation compared to PSD95. (B) Minimum Distance (End-to-End Distance): Time evolution of the end-to-end distance between the terminal residues of PSD95 (left) and PAL-PSD95 (right). The end-to-end distance of PAL-PSD95 remains consistently higher, indicating a more open conformation. (C) Root Mean Square Fluctuation (RMSF): Per-residue fluctuation profile of PSD95 (left) and PAL-PSD95 (right) calculated over the simulation trajectory. PAL-PSD95 shows markedly increased flexibility at the N-terminus and select loop regions, likely reflecting dynamic interactions with the membrane and structural rearrangements driven by palmitoylation.

**Fig. S 6:**
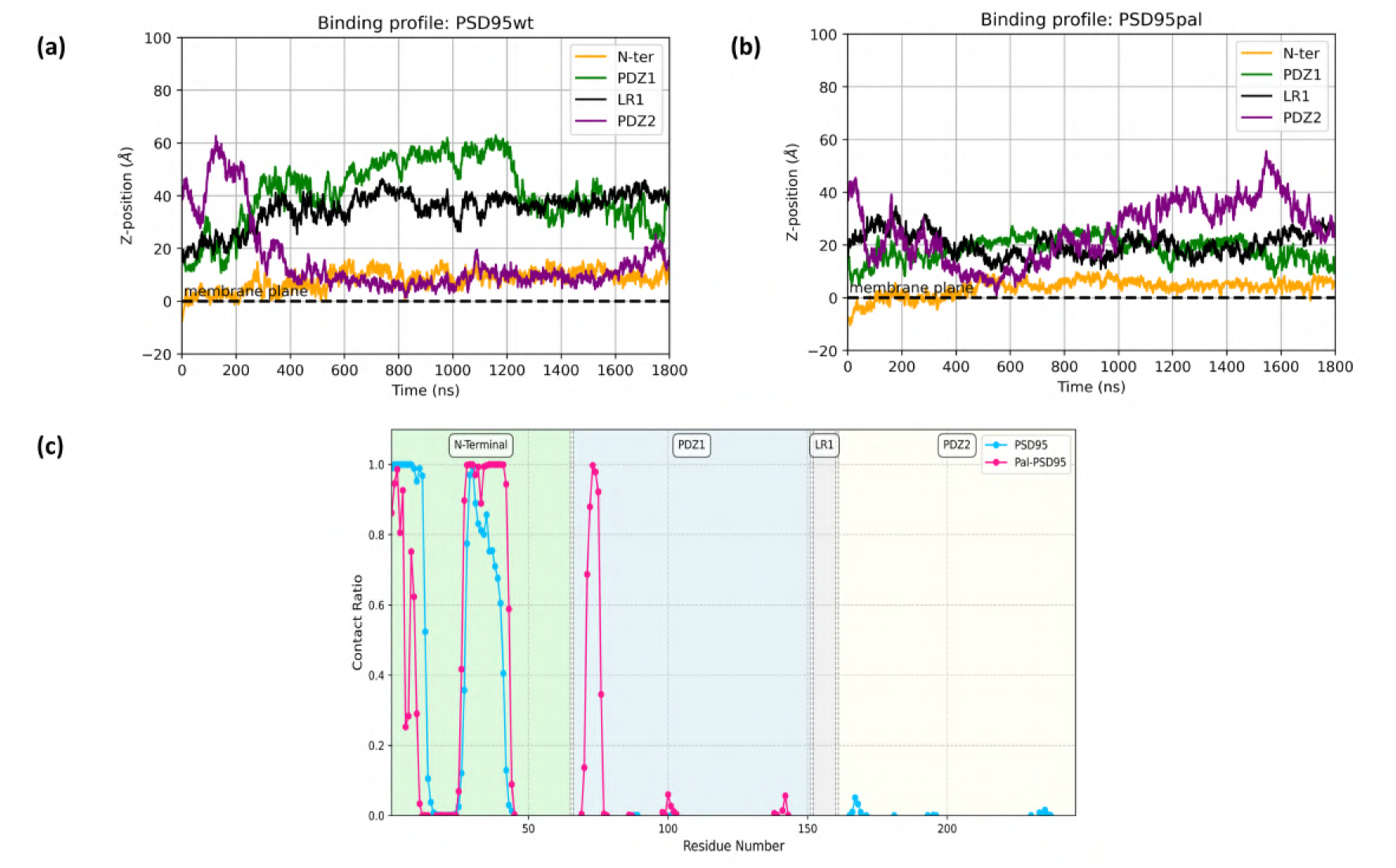
Membrane binding dynamics and lipid contact profiles of PSD-95 and palmitoylated PSD-95. (a) Z-position (Å) of different regions of PSD-95, including the N-terminal, PDZ1 with linker 1 (LR1), and PDZ2, relative to the membrane plane over time (ns) for PSD-95 (left) and palmitoylated PSD-95 (right). (b) Lipid contact ratios for each residue across PSD-95 (left) and palmitoylated PSD-95 (right). Palmitoylation enhances membrane interactions, particularly for the N-terminal and PDZ1 domains, while the PDZ2 domain shows no significant membrane contact.

## Notes

https://zenodo.org/uploads/15691830

### Competing Interest Statement

The authors have declared no competing interest.

